# Abnormal Speech Motor Control in Individuals with 16p11.2 Deletions

**DOI:** 10.1101/165316

**Authors:** Carly Demopoulos, Hardik Kothare, Danielle Mizuiri, Jennifer Henderson-Sabes, Brieana Fregeau, Jennifer Tjernagel, John Houde, Elliott H. Sherr, Srikantan S. Nagarajan

**Author notes:** Corresponding Authors: Carly Demopoulos, Ph.D., Department of Radiology & Biomedical Imaging, 513 Parnassus Avenue, S362, San Francisco, CA 94143, Srikantan S. Nagarajan, Ph.D., Department of Radiology & Biomedical Imaging, 513 Parnassus Avenue, S362, San Francisco, CA 94143.

## Abstract

Speech and motor deficits are highly prevalent (>70%) in individuals with the 600 kb BP4-BP5 16p11.2 deletion; however, the mechanisms that drive these deficits are unclear, limiting our ability to target interventions and advance treatment. This study examined fundamental aspects of speech motor control in participants with the 16p11.2 deletion. To assess capacity for control of voice, we examined how accurately and quickly subjects changed the pitch of their voice within a trial to correct for a transient perturbation of the pitch of their auditory feedback. When compared to sibling controls, 16p11.2 deletion carriers show an over-exaggerated pitch compensation response to unpredictable mid-vocalization pitch perturbations. We also examined sensorimotor adaptation of speech by assessing how subjects learned to adapt their sustained productions of formants (speech spectral peak frequencies important for vowel identity), in response to consistent changes in their auditory feedback during vowel production. Deletion carriers show reduced sensorimotor adaptation to sustained vowel identity changes in auditory feedback. These results together suggest that 16p11.2 deletion carriers have fundamental impairments in the basic mechanisms of speech motor control and these impairments may partially explain the deficits in speech and language in these individuals.

---

Speech and communication deficits can have a pervasive impact on learning, development, and quality of life and are highly prevalent in individuals with Autism Spectrum Disorders (ASD)^1,2^. There is also increasing recognition that general motor abnormalities, present in 79% of individuals with ASD^3^, predict delayed speech and language development^4–6^. Recently, given the advances in understanding the genetic basis of ASD in some individuals, an effort has been made to categorize ASD into different genetically defined subtypes^7^ by examining the behavioral, anatomical, and physiological phenotypes associated with mutations in single genes or copy number variants for contiguous genes. For example, the recurrent nearly 600 kb deletion at 16p11.2 (BP4-BP5) is one of the most common genetic etiologies of ASD and, more generally, of neurodevelopmental impairment^8–10^. Although a variety of cognitive domains have been reported to be impacted in 16p11.2 deletion carriers, the most commonly observed diagnoses are developmental coordination disorder, phonological processing disorder, language disorders, and ASD^11^. One or more speech and language diagnoses are present in 71% of all individuals with the 16p11.2 deletion^11^, including a higher than normal prevalence of childhood apraxia of speech^11–14^. These deficits seem to be specific to 16p11.2 deletion carriers, as these clinical deficits are absent in those with duplications of the same locus^13^. The etiologies of these speech production deficits, both in children with ASD and 16p11.2 deletion carriers, remain unclear, and this poses a considerable barrier to advancing treatment and intervention to improve speech in these populations.

Intact speech results from a careful orchestration of several processes to control the vocal tract^15^. For example, articulation of vowel sounds requires vocalization at the larynx while simultaneously holding the tongue and jaw in just the right shape. During articulation, we also hear the speech we are producing (auditory feedback), and if what we hear is not what we meant to say (i.e., is not consistent with our internal prediction of intended speech), we respond by correcting our speech. Thus, the control of speech requires (1) the ability to generate an expectation for the intended speech outcome (*internal speech modeling*), (2) the ability to control the muscles of the vocal apparatus to achieve that intended outcome (*vocal motor control*), and (3) the ability to accurately process and respond to auditory feedback of one's own speech in real time (*online speech error monitoring*)^15^. How these specific fundamental processes of speech motor control are impacted in children with ASD and the 16p11.2 deletion is poorly understood; however, prior studies examining low level speech motor control in children with apraxia of speech and other speech sound disorders suggest that these speech disorders may result from impairment in internal speech modeling, requiring a compensatory increased reliance on auditory feedback for speech production^16–19^. Given the highly penetrant speech production deficits in children with the 16p11.2 deletion, examination of these low level speech motor control processes in this group offers an opportunity to identify discrete points of failure in the speech production system. In particular, better understanding the role of auditory feedback in the control of speech in these participants could lead to insights into potential targets for speech and language intervention and directions for the development of novel therapeutics.

Thus, the present study investigates the speech motor control and auditory feedback processing during speech in participants with the 16p11.2 deletion. In particular, we assessed (1) how quickly participants reacted to changes in auditory feedback, and (2) how well they could learn from their auditory feedback. We made these assessments by examining two well-studied phenomena: (1) The *pitch perturbation reflex* is seen when the pitch of speakers’ auditory feedback of their ongoing speech is suddenly and unexpectedly perturbed^20^. Speakers quickly respond to such perturbations (within 100-200ms) by compensating (i.e., changing their pitch to negate the perceived effect of the pitch perturbation). The pitch perturbation reflex thus measures how speakers immediately react to auditory feedback for online speech error monitoring and correction. (2) *Speech sensorimotor adaptation* can be seen when the formants of speakers’ auditory feedback are consistently altered on repeated vowel productions^21,22^. Such formant alterations change how speakers perceive their vowel sounds: the vowel they hear in their auditory feedback does not match the vowel they intended to produce. Speakers respond to this mismatch by adapting their vowel productions over repeated trials, with each successive production changed to more fully counter the perceived effects of the formant alterations. Speech sensorimotor adaptation thus measures how well speakers can use auditory feedback to incorporate long-term changes in their speech production. Both the pitch perturbation reflex and speech sensorimotor adaptation response have proven to be sensitive indicators of abnormal speech feedback processing in other neurological conditions^23–25^, and were therefore natural choices for assessing feedback processing in our participants.

## Methods

### Participants

Twelve 16p11.2 deletion carriers (three females and nine males, mean age=12.4, SD=3.2, range=8.3-18.3) and six unaffected siblings (one female and 5 males, mean age=12.8, SD=1.5, range=11.3-15.4) were enrolled from the 2015 Simon's Variation in Individuals Project Family Meeting in Falls Church, VA. The study was offered to attendees as a speech and hearing study. In addition, five neurotypical control participants (two females and three males, mean age=12.4, SD=1.8, range=10.3-14.2) were recruited through community advertisement. Subjects were matched for age across the groups - deletion carriers, unaffected sibling controls and neurotypical controls, F(2,20)=.037, p=.96. Inclusion criteria were (1) presence of a BP4-BP5 16p11.2 deletion, sibling of a participant with a BP4-BP5 16p11.2 deletion who did not have the 16p11.2 deletion or any other known genetic or neuropsychiatric disorder, (2) age range of 8-18 years, and (3) ability to produce a sustained phonation (e.g., say “ah” for 2-3 seconds). Informed consent and assent were obtained and all procedures were approved by and were performed in accordance with the requirements of the Committee on Human Research at University of California-San Francisco.

### Measures

Participants were assessed on two measures of sensitivity to auditory feedback during speech: a Pitch Perturbation Task and a Formant Adaptation Task. For both of these tasks the auditory feedback the participant hears while speaking is altered in real time. To identify potential confounds related to hearing abilities, auditory brainstem responses (ABRs) were also assessed for participants with the 16p11.2 deletion and their siblings. In the 16p11.2 deletion group, time constraints prevented two participants from completing the ABR task and three from completing the speech production tasks. Additionally, one participant discontinued participation in the study due to extreme sound sensitivity to the ABR task stimuli. In the neurotypical control group, one participant was excluded from the speech adaptation task due to technical issues.

#### Pitch Perturbation Task

This task measured subjects’ immediate corrective responses to unexpected perturbations of the pitch of the auditory feedback of their ongoing speech. In each trial, participants produced and sustained the vowel sound, “ah”, for at least 2.5 seconds into a microphone. Speech from the microphone was sent through a digital signal processing (DSP) system before being returned to the subjects’ headphones as auditory feedback of their speech. The DSP was capable of altering speech feedback (in either pitch or formant frequencies) in real time (12ms feedback delay)^26–28^. During each trial the participant's auditory feedback was randomly raised or lowered in pitch by 1/12^th^ of an octave (+/-100 cents) at a variable lag from speech onset for 400ms duration (Figure 1). This process was repeated for 74 trials with an intertrial interval of 2.5 seconds. This task measures how quickly participants adjust their speech within each trial in order to compensate for the perceived speech error created by the mid-vocalization pitch perturbation. The compensation is subtle, and the participants may not be aware of it. Peak magnitude of compensation is determined by peak response change from pre-to post-perturbation baseline by: cents change = 100 × [12 × log2(pitch response peak (Hz) / mean pitch frequency of pre-perturbation baseline (Hz))]^27^. Baseline begins at voice onset and ends at onset of perturbation. For statistical comparison between groups, the mean of the peak compensation for each subject was calculated by extracting the peak deviation from baseline pitch in each trial and computing the average of the peak compensation response across trials. Peak latency from perturbation onset was also computed.

**Figure 1.**
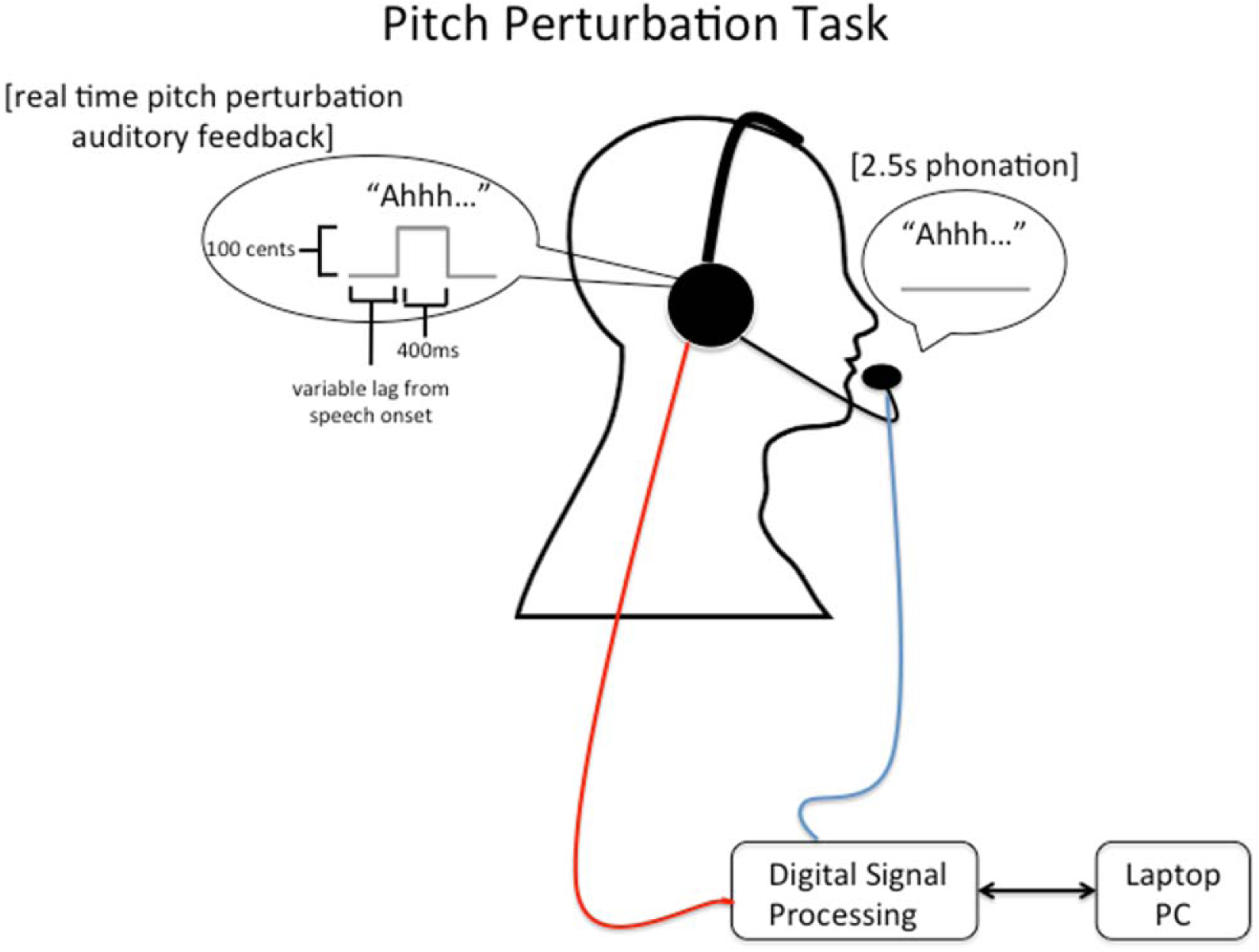
Apparatus for Pitch Perturbation Task. The participant phonates the vowel sound “ahh” into a headset microphone for the duration of a visual cue presented for 2.5 seconds. The microphone signal is then digitally processed to randomly shift the pitch of the recorded speech at a variable lag from speech onset up or down by 100 cents in real time creating an effect of altered auditory feedback in the participants headphones.

#### Formant Adaptation Task

This task measured how much subjects learned to adjust their speech production to consistently minimize perceived speech errors (altered formants) in their auditory feedback. The microphone/DSP/headphone setup was the same as for the pitch perturbation task. In each trial, participants said the word “bed” upon cue into a microphone, and heard auditory feedback of their production in the headphones, with an inter-trial interval of 2.5 seconds. For the first 20 trials, the DSP provided subjects with unaltered feedback of their productions. In the following 30 trials, however, the DSP altered the formants (F1 and F2) of the auditory feedback so that subjects heard the vowel /ae/ when they produced the vowel /eh/. F1 was raised by 200 Hz and F2 was reduced by −250 Hz during the altered block of 30 trials. Thus, on these trials, when subjects produced the word “bed” they heard the word “bad” (Figure 2). To compute adaptation we first represented the applied formant shift and produced formants as vectors from baseline formant values in an F1-F2 (first and second formants) Euclidean space. Produced formant frequencies were extracted from spectrograms of entire vowel utterances. Formant dropouts and instances of incorrect tracking were excluded from the analysis. We then determined the scalar projection of the produced formants vector on the formant shift vector. Percentage adaptation was calculated as −100 x (scalar projection) / (magnitude of applied shift). aloud into a headset microphone. The digital signal processor then alters the formants (F1 and F2) of the auditory feedback so that participants hear the vowel /ae/ when they produced the vowel /eh/. F1 was raised by 200 Hz and F2 was reduced by −250 Hz during the altered block of 30 trials. Thus, on these trials, when subjects produced the word “bed” they heard the word “bad”.

**Figure 2.**
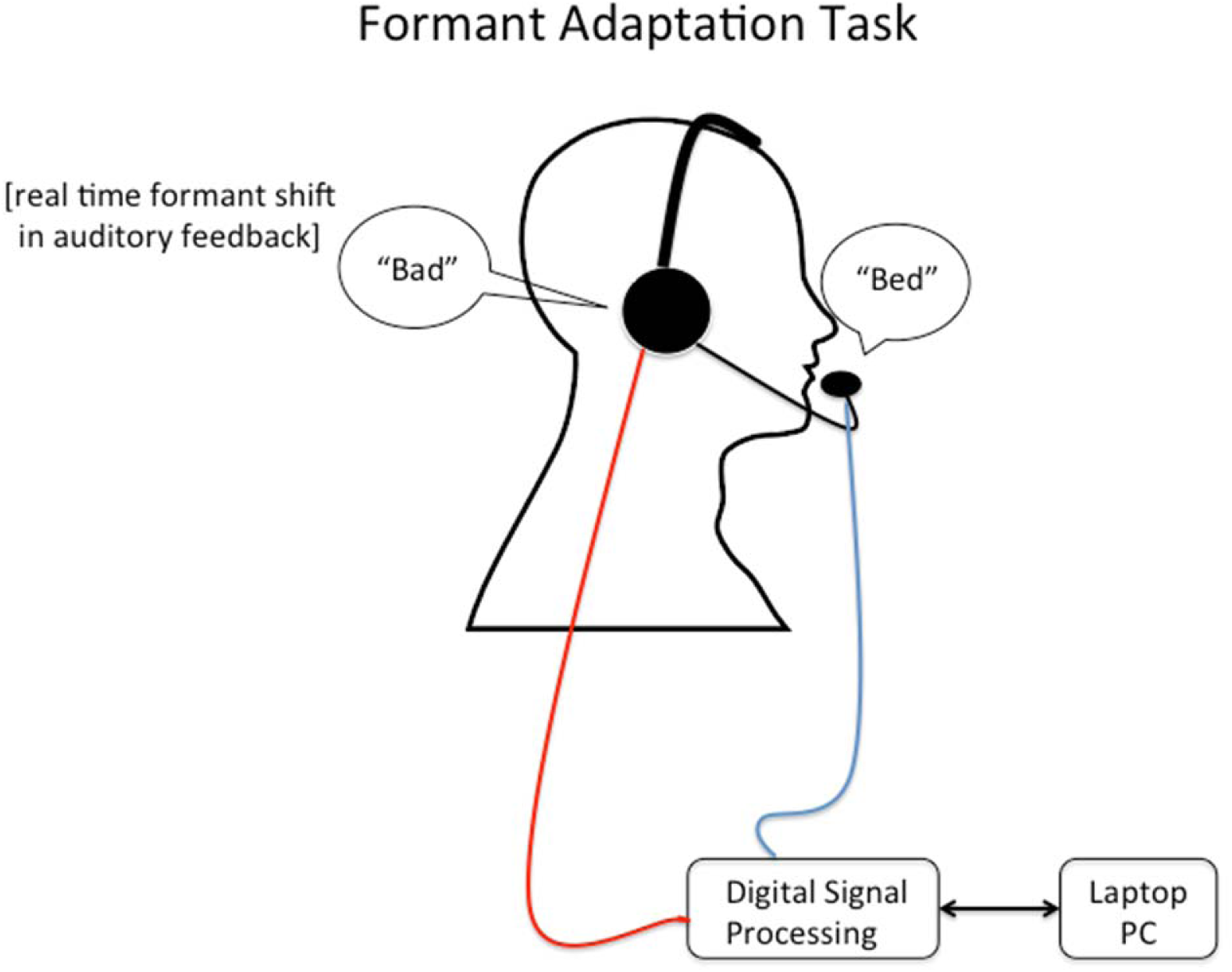
Apparatus for Formant Adaptation Task. The participant is cued to read the word “bed”

#### Auditory Brainstem Response (ABR) Task

Because the goal of the study was to examine the role of auditory feedback in speech for a group of participants who may have abnormalities in auditory processing, we also administered an auditory brainstem response task to assess basic auditory processing in the absence of speech. This task required the participants to passively listen to clicking sounds through headphones while auditory brainstem responses were collected through electrodes placed on both earlobes with a reference electrode on the forehead. The ABR is a neurophysiological response reflecting activity in peripheral aspects of the auditory system (e.g., cochlear nucleus, medial lemniscus, and brainstem nuclei). Because this is measured passively, ABRs allow for estimation of pronounced hearing loss without the cooperation and cognitive task demands required by behavioral methods for estimating hearing thresholds; however, the ABR is not a sensitive measure of mild hearing loss, and it generally does not provide information on potential hearing loss at low frequencies. Data were collected in response to 75 dB clicks presented at a rate of 11.1 Hz, with a total of 1000 stimuli presented to each ear. Data were analyzed by a licensed clinical audiologist (JHS) with respect to the latencies of ABR waves I, III, and V, wave I–III and III–V interpeak latencies, the wave V amplitude and interaural amplitude differences, and overall morphology.

#### Hypotheses and Data Analytic Plan

Based on prior literature documenting speech abnormalities in children and adolescents with 16p11.2 deletions^12^, we hypothesized that deletion carriers would show abnormalities on the pitch perturbation and formant adaptation tasks. To test this hypothesis we performed three independent samples t-tests to examine group differences in (1) peak magnitude and (2) latency of compensation (assessed via the pitch perturbation task) and (2) adaptation of the internal speech model (assessed via the formant adaptation task). Independent samples t-tests were also performed on ABR data to assess group differences in wave V latencies, amplitude, and interaural amplitude differences to ensure that group differences in compensation or adaptation were not an artifact of gross hearing impairment in16p11.2 deletion carriers.

Eight participants in the 16p11.2 deletion group and eleven participants in the control group completed the pitch perturbation task; however, data from one participant in the 16p11.2 deletion group was excluded from analyses due to excessive baseline noise. Further, two sibling control participants did not produce a compensation response and instead adjusted their speech in the same direction as the speech perturbation (“following” response). Because this following response is thought to be categorically different from a compensation response, and because it had the potential to artificially inflate group differences, compensation data from these two participants were also excluded from analyses, resulting in a comparison of seven participants with 16p11.2 deletions and nine control participants.

Finally, to determine whether adaptation responses were due to learning across trials (as opposed to within-trial compensation in response to the feedback alteration), a repeated-measures analysis of variance was performed. Between-subjects, within-subjects, and interaction effects of adaptation across the chronology of trials (binned into group of 5 trials) were examined for the formant adaptation task to determine if adaptation increased over the course of trials and if this increase differed between groups.

## Results

### Pitch Perturbation Task

The average compensatory response to the applied pitch perturbation, over the pre-perturbation through post-perturbation vocalization interval, is shown across subjects for the deletion carriers and their siblings in Figure 3. Vocalizations were similar between groups at baseline and until 200ms following the onset of the feedback perturbation of pitch. However, the groups’ pitch compensation responses subsequently diverged in their vocal response to the auditory feedback perturbation, wherein the deletion carriers show an enhanced compensatory response when compared with controls starting at around 200ms post-perturbation onset and persisting through the response interval. The mean percent compensation response to pitch feedback perturbations was significantly enhanced in 16p11.2 deletion carriers (47.50 +/-28.77%) when compared to controls (21.79 +/-11.42%), t(14)=-2.46, p=.027. There were no significant differences between groups for peak latency of compensation response for 16p11.2 deletion carriers (.66 +/-.20s) and control participants (.59 +/-.17s), t(14)=.813, p=.43. Individual participant compensation responses are illustrated in Figure 4.

**Figure 3.**
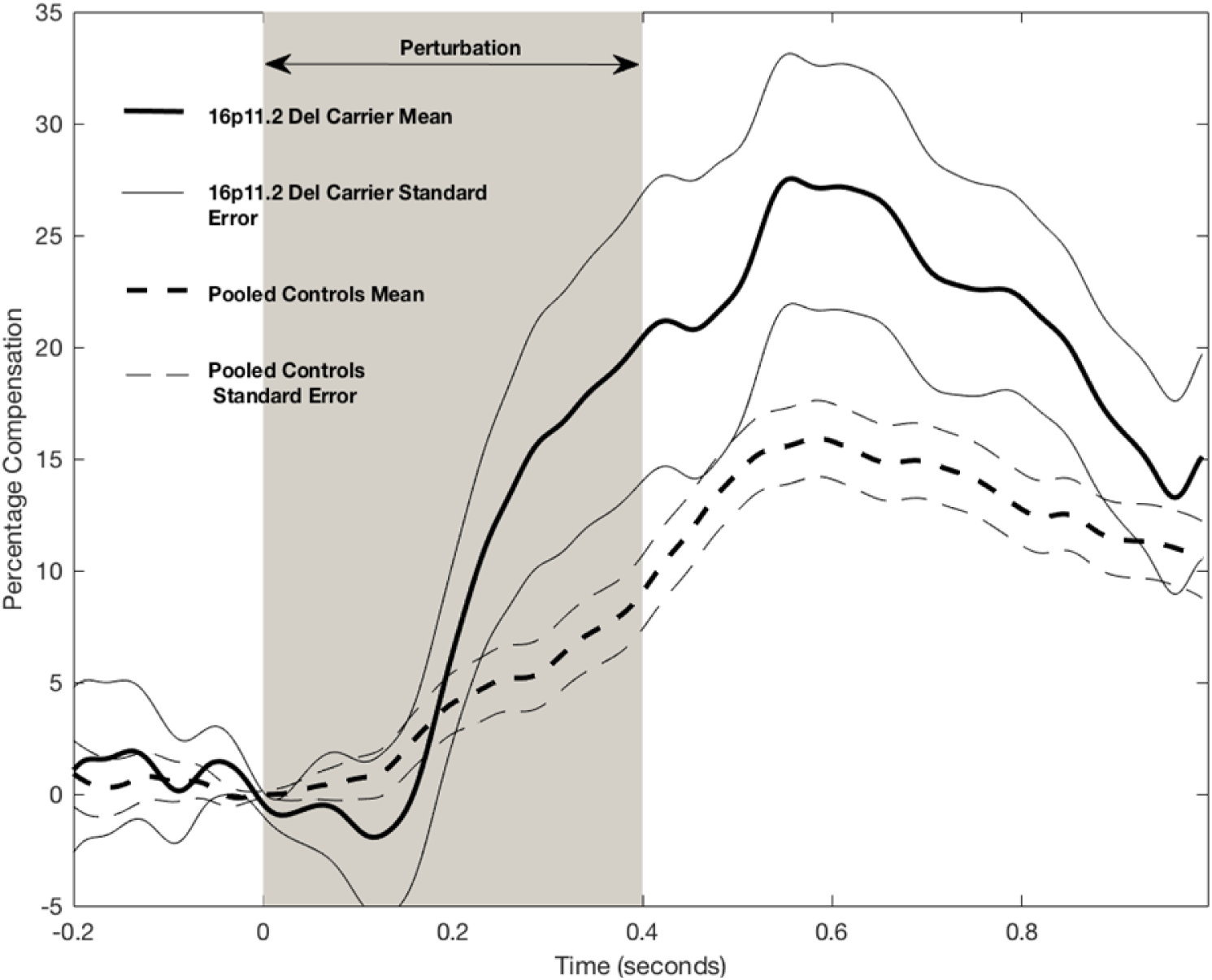
Average changes in compensation over the course of the vocalization in the Pitch Perturbation Task for participants with 16p11.2 deletions and the control group. The shaded area indicates the onset of auditory feedback perturbation. 16p11.2 deletion carriers demonstrate an exaggerated compensation response relative to the control participants.

**Figure 4.**
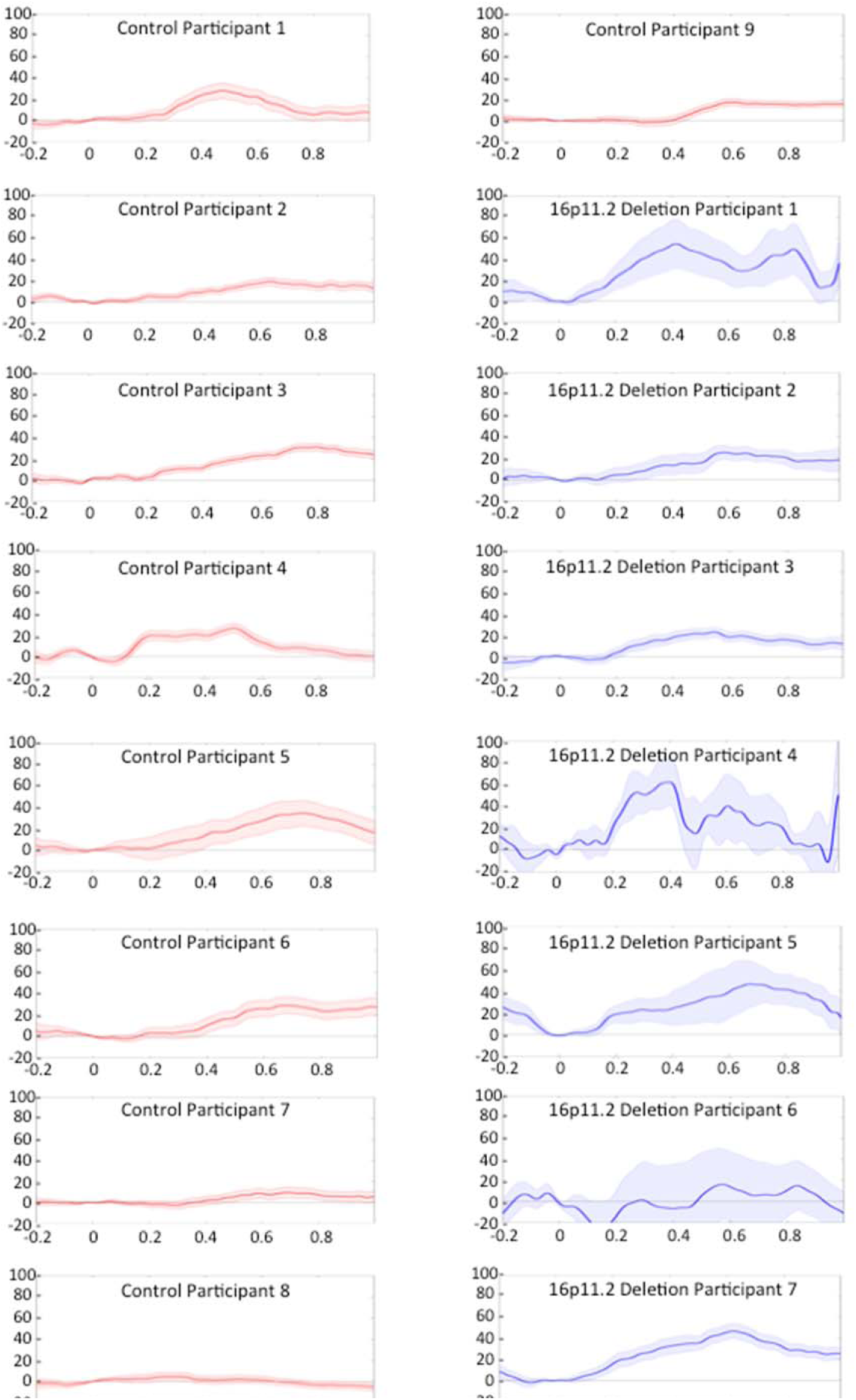
Individual participant mean responses to pitch perturbation task. Mean percentage compensation (vertical axis) responses (solid lines) and error bars (shaded lines) are depicted for participants in the 16p11.2 deletion (blue) and control (red) groups across time from perturbation onset in seconds (horizontal axis).

### Formant Adaptation Task

We examined the magnitude of formant adaptation in response to repeated perturbations of formants in auditory feedback in the eight participants in the 16p11.2 deletion group and ten participants in the control group who completed the speech adaptation task. While control participants learned to adjust their speech in anticipation of the altered auditory feedback, the 16p11.2 deletion carriers did not adjust their speech in response to predictable auditory feedback alterations. Figure 5 illustrates the mean group differences in formant adaptation response across 16p11.2 deletion carriers and control participants. In contrast to within trial pitch compensation responses, reduction in the magnitude of formant adaptation in response to repeated perturbation of formants in auditory feedback across trials was also significant in 16p11.2 deletion carriers (4.22 +/-5.17%) when compared to the control group (13.2 +/-5.01%), t(16) = −3.72, p=0.002.

**Figure 5.**
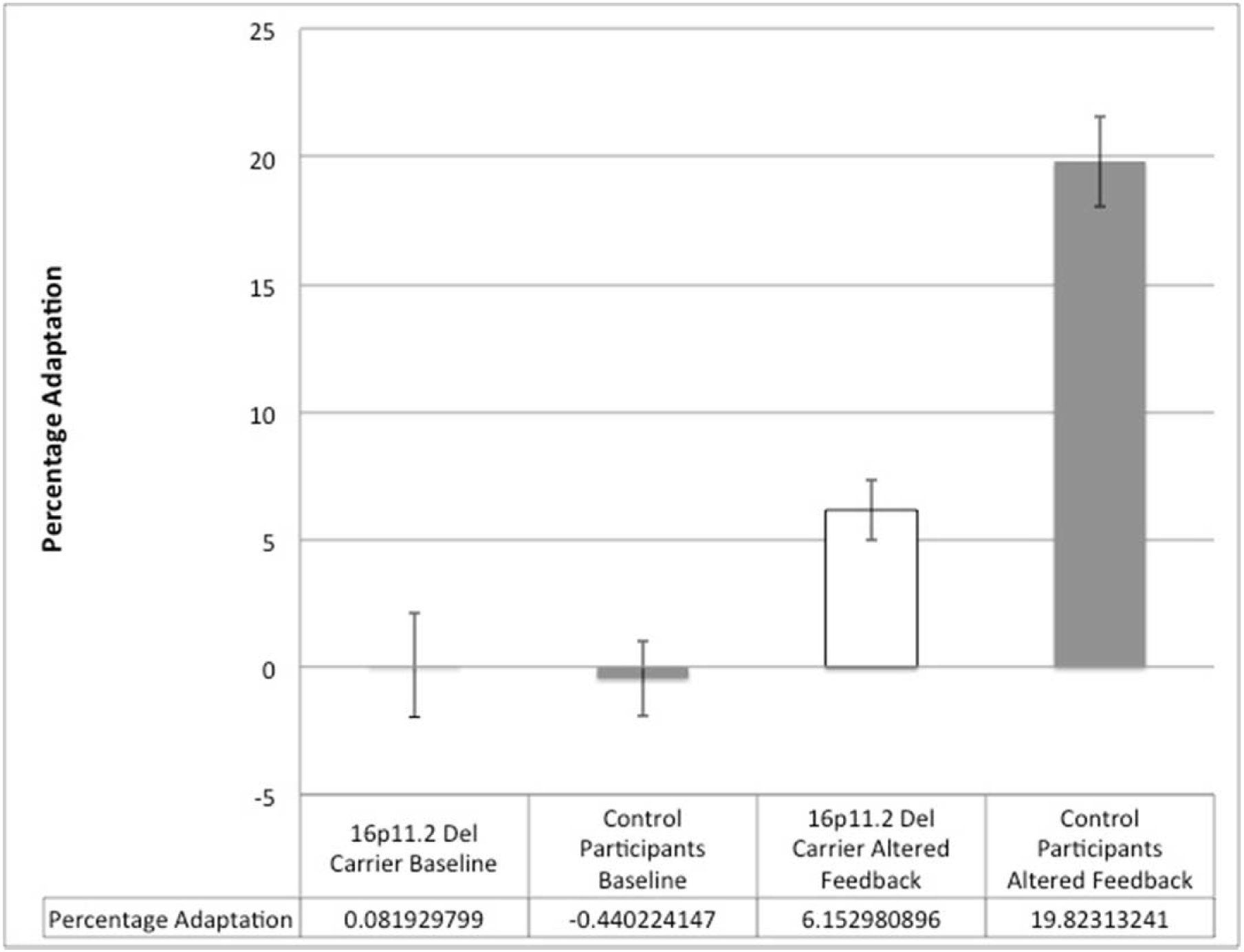
Mean group adaptation across trials for the baseline and altered feedback conditions in the Formant Adaptation Task. While groups did not differ at baseline, significant group differences were identified in the altered feedback condition, with the deletion carriers showing significantly reduced adaptation.

Next, to determine whether the adaptation response demonstrated by participants in the control group was actually due to learning across trials (as opposed to within-trial compensation in response to the feedback alteration), a repeated-measures analysis of variance was performed. Between-subjects, within-subjects, and interaction effects of adaptation across the chronology of trials (binned into group of 5 trials) were examined for the formant adaptation task. Results of between-subjects analyses indicated that the control group demonstrated significantly greater adaptation for altered feedback trials then the 16p11.2 deletion group, F(1,16)=6.59, p=.021, partial η^2^=.292. Within-subject effects were also statistically significant, F(9,144)=5.81, p<.001, partial η^2^=.266. Group by trial interaction effects were not statistically significant F(9,144)=1.77, p=.079, partial η^2^=.099; however, linear contrasts were significant for both within-subjects, F(1,16)=44.31, p<.001, partial η^2^=.735, and interaction effects, F(1,16)=11.51, p=.004, partial η^2^=.418. Figure 6 depicts these linear effects, as participants in the control group demonstrated greater learning across trials than participants with 16p11.2 deletions. These findings suggest that increases in percent compensation across trials for the control group was a result of learning over the course of the altered feedback trials and not an artifact of within-trial compensation.

**Figure 6.**
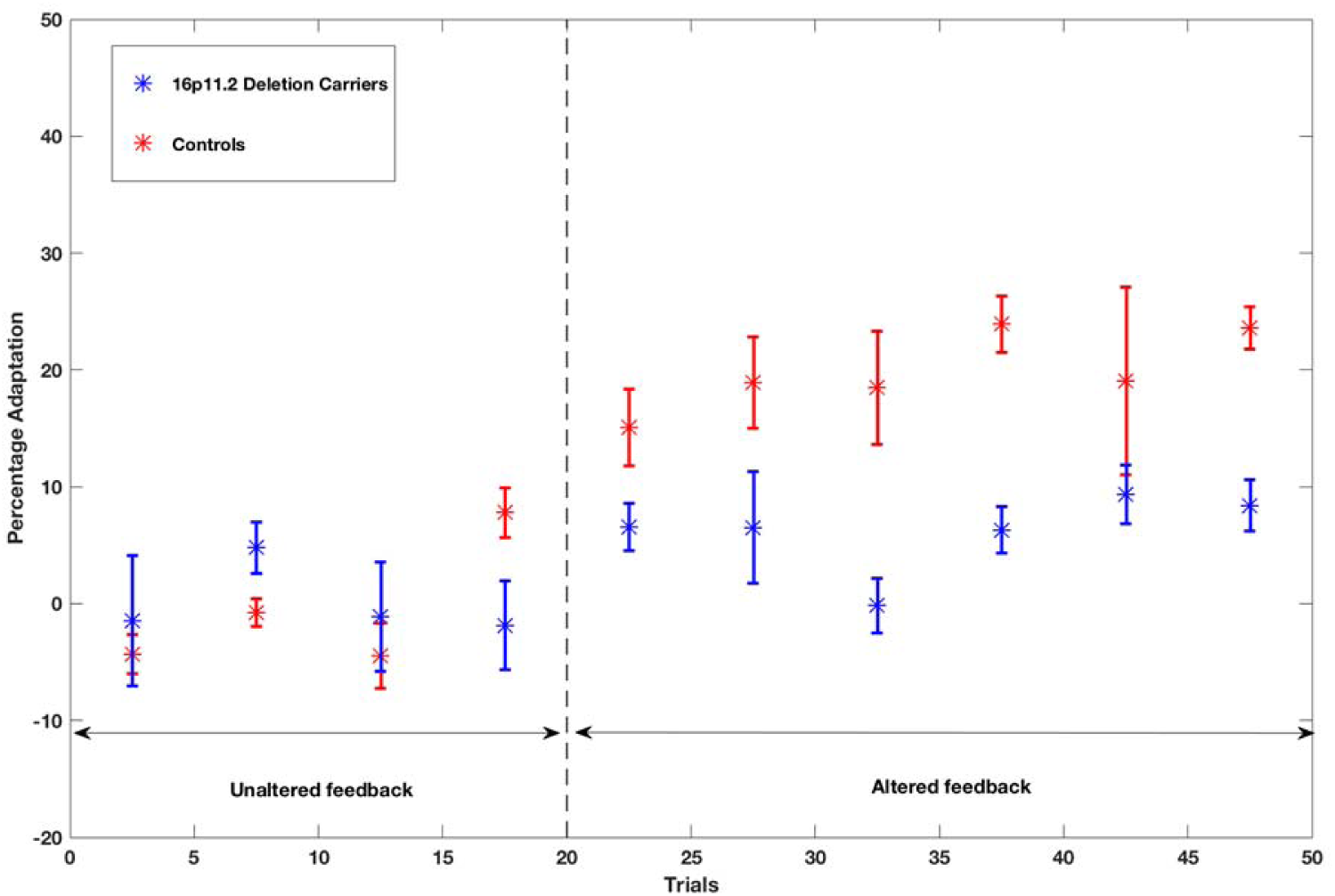
Mean adaptation responses across trials grouped into bins of five trials. Significant linear effects of group and group by trial number interaction are illustrated. Following the baseline trials in which feedback was unaltered (trials 1-20), the control group demonstrated a gradual increase in adaptation with increasing number of altered feedback trials (21-50), whereas the 16p11.2 deletion group show a weaker adaptation response.

### ABR Task

Finally, ABR data were evaluated in the nine 16p11.2 deletion carriers and five sibling controls who completed the task to assess for evidence of severe hearing impairment and Wave V abnormalities. No evidence of hearing impairment was identified by our licensed clinical audiologist (JHS) for any of the participants, and no group differences were detected for independent samples t-tests examining differences in Wave V latency or amplitude variables (Table 1), indicating that it is unlikely that group differences in speech psychophysics could be an artifact of hearing impairment in our 16p11.2 deletion group.

**Table 1.**
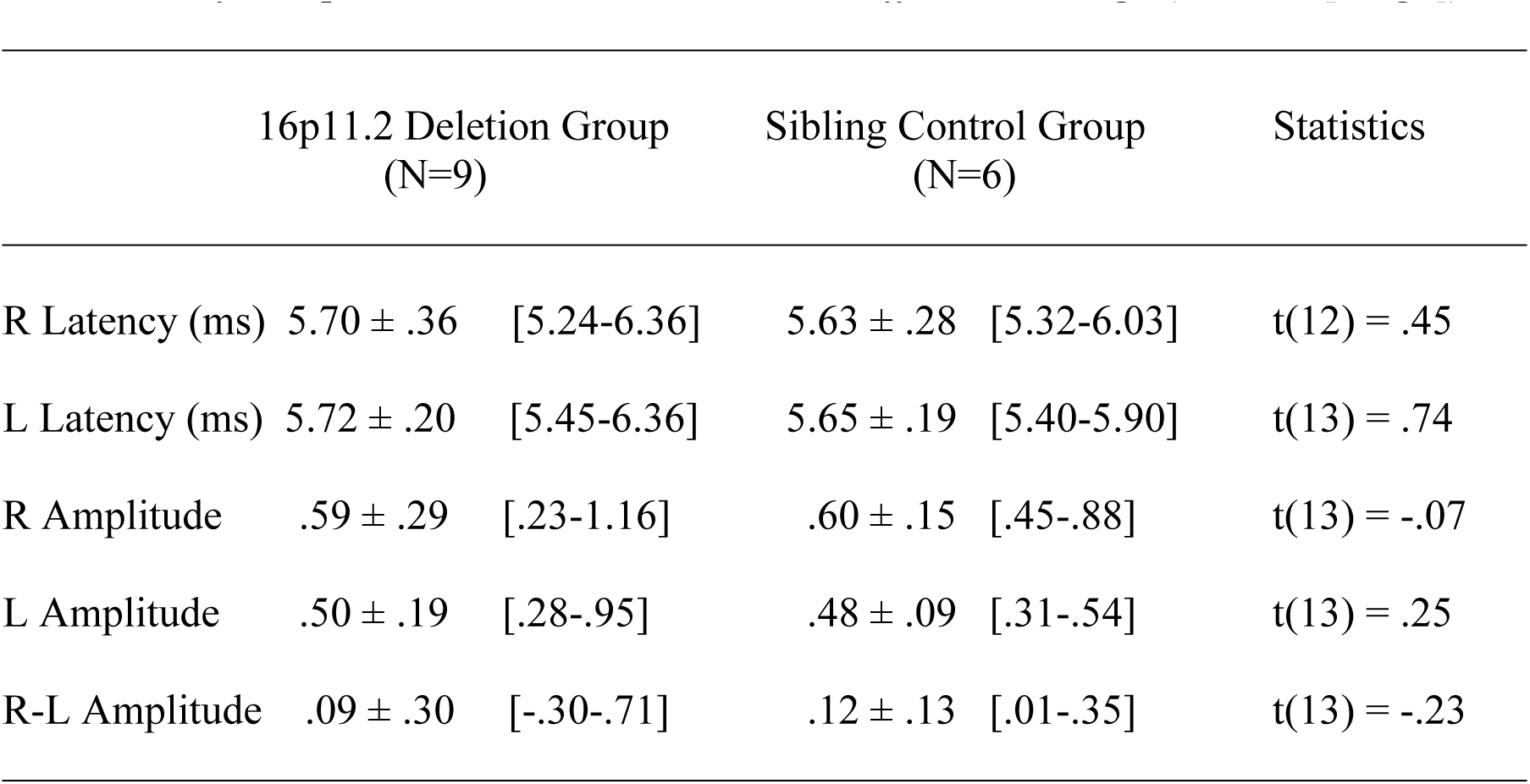
*ABR Results for 16p11.2 Deletion Carriers and Unaffected Siblings* (M ± SD [*range*])

## Discussion

This study examined fundamental processes of speech motor control in participants with 16p11.2 deletions. Specifically, we examined sensitivity to auditory feedback processing during speech production using measures of (1) online error monitoring and resultant compensatory speech in response to mid-utterance auditory feedback alterations, and (2) across-trial adaptation of vowel production in response to computer generated consistent auditory feedback errors. First, our results support the hypothesis that 16p11.2 deletion carriers show abnormalities in compensation, as carriers demonstrated an overcompensation response to pitch perturbations. Our findings extend and confirm hints of overcompensation reported in some individuals with ASD, where a subset of participants with idiopathic autism with weaker language ability showed increased pitch compensation^29^. One possible explanation for this overcompensation response is that auditory processing dysfunction results in distorted perception of the mismatch between expected and actual auditory feedback, thereby driving overcompensation by way of inaccurate perception of the target during speech correction. However, similar timing of the peak of the compensatory responses suggests that duration of auditory error processing is not impaired in 16p11.2 deletion carriers. A second possibility is the heightening of the motoric responses to auditory feedback errors in 16p11.2 deletion carriers.

Our data also showed reduced speech model adaptation following consecutive trials in which auditory feedback of vowel sounds was consistently altered in a predictable way. A reduced adaptation response has been reported in typically developing younger children and to an even greater extent in toddlers^30^, which may suggest a maturational dysfunction in one or more speech production mechanisms in 16p11.2 deletion carriers. This reduced speech model adaptation when taken together with the strong compensation response during the pitch perturbation task suggests that while deletion carriers were able to detect and make some effort to correct perceived speech errors in real time, they were impaired in learning to anticipate the effects of the highly predictable alterations in auditory feedback in their vowel productions.

There are several possible interpretations for these results. Normal speakers are assumed to have internal pre-stored representations of the motor programs they need for producing the repertoire of speech sounds corresponding to the languages in which they converse. These internal representations allow speakers to correctly produce speech even in the absence of sensory feedback; they need not rely on immediate sensory feedback for the moment-to-moment control of their speech. Nevertheless, sensory feedback—and especially auditory feedback—plays a critical role in maintaining the accuracy of these internal representations, which are constantly being adjusted to eliminate any consistent sensory feedback prediction errors^15^. In comparison, participants with 16p11.2 deletions may have impairments in sensorimotor predicting/learning, resulting in an inability to anticipate the consistent auditory feedback prediction errors created by the formant feedback alterations. Alternatively, they may not have any reliable pre-stored representations to govern their speech production, and may instead be forced to rely on immediate sensory feedback for the moment-to-moment control of their speech. In the context of intact ability to generate internal speech motor programs, they may only be able to adapt them in a rigid way, perhaps due to a limited repertoire of speech motor programs, or rote approach to speech learning, resulting in difficulty in flexibly adapting the internal representations of their speech motor programs. Difficulty with generating or adapting these internal speech motor representations could result in a compensatory over-reliance on feedback for error correction during speech and subsequently, the reduced adaptation and overcompensation response observed in our sample. Over-reliance on auditory feedback has been interpreted as a compensatory response to weak feedforward control of speech in previous studies of adults with Parkinson's Disease^23,31^ and children with apraxia of speech (AOS)^16,17^. Terband et al. (2014) examined a broader group of children with speech sound disorders (not limited to AOS) and found a failure to effectively adapt to altered speech feedback^19^, which was strongly associated with poor performance on a nonword repetition task. Computational models have also been used to demonstrate that an increased reliance on feedback from speech can account for several characteristics of AOS^18^.

### Limitations And Future Directions

Several limitations of the present study must be acknowledged. Most notably, the small sample size resulted in limited power to detect modest effects and greater potential for spurious findings. Therefore, these findings should be replicated with a larger sample. A second limitation was our inability to examine post-feedback alteration responses for the formant adaptation task. While we were able to demonstrate that participants in the control group did increase their adaptation responses gradually over the course of altered-feedback trials, suggesting that learning had taken place, a post-feedback phase would have directly assessed for evidence of learning. In addition, our compensation and adaptation tasks employed different types of feedback alterations (i.e., altered pitch for compensation and altered formants for adaptation). While these findings provide some preliminary evidence that there is an overreliance on auditory feedback and a weakness in feedforward modeling for individuals with 16p11.2 deletions, these paradigms will need to be administered with the same auditory feedback across compensation and adaptation tasks in order to draw definitive conclusions, as prior research has demonstrated a dissociation between the compensation response in pitch versus formant feedback alterations in other clinical populations, such as Parkinson's Disease^32^.

Future studies should include a more comprehensive assessment of auditory processing, speech, and language function, along with assessment of vocal motor control to fully understand the impact of sensory and motor function on abnormal speech perturbation responses. Future studies that combine speech psychophysics with neuroimaging are necessary to determine (1) whether these abnormal compensation and adaptation responses are associated with difficulty in generating or adapting internal speech representations, (2) whether such difficulties result in a compensatory overreliance on auditory feedback for the moment to moment control of speech, and (3) whether observed speech motor control abnormalities are directly associated with speech and language deficits.

## Conclusions

Intact speech results from a careful orchestration of several fundamental processes of speech motor control, and dysfunction in any part of this system could impact speech production in numerous ways. If we are to understand the causal factors in the emergence of speech deficits in 16p11.2 deletion carriers, we must first understand these fundamental processes associated with the earliest presenting symptoms, such as sensory and motor dysfunction, and their subsequent impact on more complex processes, such as speech and language. This study suggests some initial directions for this line of inquiry.

## Acknowledgments

This project was funded in part by the Simons VIP Project grant and National Institutes of Health grants (R01DC004855, R01DC010145, R21NS076171 and R01DC013979), National Science Foundation grant (BCS 1262297), and the US Department of Defense grant (W81XWH-13-1-0494). We are grateful to all of the families at the participating 2015 Simon's Variation in Individuals Project Family Meeting in Falls Church, VA, as well as the Simons VIP Consortium. We would also like to thank Wendy Chung for her thoughtful comments that greatly improved the manuscript.

## Author Contributions

CD was responsible for data collection, processing, analysis, interpretation and writing of the manuscript. HK was responsible for data collection, processing, and analysis. DM was responsible for data collection. JHS was responsible for data processing. BF was involved in study coordination, consenting and data management. JT was involved in study execution and coordination of the data collection meeting. JH was responsible for task development and was involved in manuscript review. EHS and SSN were involved in conceptualization and design of the project and writing of the manuscript.

## Additional Information

The author(s) declare no competing financial interests.

## References

1. Tager-Flusberg, H., Paul, R. & Lord, C. in Handbook of Autism and Pervasive Developmental Disorders, 3rd Edition (eds. Volkmar, F. R., Paul, R. & Klin, A.) 335–364 (2005).

2. Van Agt, H., Verhoeven, L., Van Den Brink, G. & De Koning, H. The impact on socio-emotional development and quality of life of language impairment in 8-year-old children. Dev. Med. Child Neurol. 53, 81–88 (2011).

3. Lai, M.-C., Lombardo, M. V & Baron-Cohen, S. Autism. Lancet 383, 896–910 (2014).

4. Gernsbacher, M. A., Sauer, E. A., Geye, H. M., Schweigert, E. K. & Goldsmith, H. Infant and toddler oral-and manual-motor skills predict later speech fluency in autism. J. Child Psychol. Psychiatry Allied Discip. 49, 43–50 (2008).

5. Bhat, A. N., Galloway, J. C. & Landa, R. J. Relation between early motor delay and later communication delay in infants at risk for autism. Infant Behav. Dev. 35, 838–846 (2012).

6. Sacrey, L.-A. R., Germani, T., Bryson, S. E. & Zwaigenbaum, L. Reaching and grasping in autism spectrum disorder: a review of recent literature. Front. Neurol. 5, 6 (2014).

7. Simons, T. et al. Simons Variation in Individuals Project (Simons VIP): A genetics-first approach to studying autism spectrum and related neurodevelopmental disorders NeuroView. Neuron 73, 1063–1067 (2012).

8. Kumar, R. A. et al. Recurrent 16p11.2 microdeletions in autism. Hum. Mol. Genet. 17, 628–638 (2008).

9. Weiss, L. A. et al. Association between microdeletion and microduplication at 16p11.2 and autism. N. Engl. J. Med. 358, 667–675 (2008).

10. Zufferey, F. et al. A 600 kb deletion syndrome at 16p11.2 leads to energy imbalance and neuropsychiatric disorders. J. Med. Genet. 49, 660–668 (2012).

11. Hanson, E. et al. The cognitive and behavioral phenotype of the 16p11.2 deletion in a clinically ascertained population. Biol. Psychiatry 77, 785–793 (2015).

12. Fedorenko, E. et al. A highly penetrant form of childhood apraxia of speech due to deletion of 16p11.2. Eur. J. Hum. Genet. 24, 302–306 (2016).

13. Hippolyte, L. et al. The number of genomic copies at the 16p11.2 locus modulates language, verbal memory, and inhibition. Biol. Psychiatry 80, 129–139 (2016).

14. Shinawi, M. et al. Recurrent reciprocal 16p11.2 rearrangements associated with global developmental delay, behavioural problems, dysmorphism, epilepsy, and abnormal head size. J. Med. Genet. 47, 332–341 (2010).

15. Houde, J. F. & Nagarajan, S. S. Speech Production as State Feedback Control. Front. Hum. Neurosci. 5, 1–14 (2011).

16. Maas, E., Mailend, M.-L. & Guenther, F. H. Feedforward and feedback control in apraxia of speech: Effects of noise masking on vowel production. J. Speech, Lang. Hear. Res. 58, 185–200 (2015).

17. Iuzzini-Seigel, J., Hogan, T. P., Guarino, A. J. & Green, J. R. Reliance on auditory feedback in children with childhood apraxia of speech. J. Commun. Disord. 54, 32–42 (2015).

18. Terband, H., Maassen, B., Guenther, F. H. & Brumberg, J. Computational neural modeling of speech motor control in childhood apraxia of speech (CAS). J. Speech, Lang. Hear. Res. 52, 1595–1609 (2009).

19. Terband, H., van Brenk, F. & van Doornik-van der Zee, A. Auditory feedback perturbation in children with developmental speech sound disorders. J. Commun. Disord. 51, 64–77 (2014).

20. Burnett, T. A., Freedland, M., Larson, C. & Hain, T. Voice F0 responses to manipulations in pitch feedback. J. Acoust. Soc. Am. 103, 3153–61 (1998).

21. Houde, J. F. & Jordan, M. I. Sensorimotor adaptation in speech production. Science (80-.). 279, 1213–6 (1998).

22. Houde, J. F. & Jordan, M. I. Sensorimotor adaptation of speech: Compensation and adaptation. J. Speech, Lang. Hear. Res. 45, 295–310 (2002).

23. Mollaei, F., Shiller, D. M. & Gracco, V. L. Sensorimotor adaptation of speech in Parkinson's Disease. Mov. Disord. 28, 1668–1674 (2013).

24. Daliri, A., Wieland, E. A., Cai, S., Guenther, F. H. & Chang, S. Motor adaptation is reduced in adults who stutter but not in children who stutter. Dev. Sci. (2017). doi:10.1111/desc.12521

25. van den Bunt, M. et al. Increased response to altered auditory feedback in dyslexia: A weaker sensorimotor magnet implied in the phonological deficit. J. Speech, Lang. Hear. Res. 60, 654–667 (2017).

26. Katseff, S., Houde, J. & Johnson, K. Partial compensation for altered auditory feedback: A tradeoff with somatosensory feedback ? Lang. Speech 55, 295–308 (2012).

27. Kort, N. S., Nagarajan, S. S. & Houde, J. F. A bilateral cortical network responds to pitch perturbations in speech feedback. Neuroimage 86, 525–535 (2014).

28. Ranasinghe, K. G. et al. Abnormal vocal behavior predicts executive and memory deficits in Alzheimer's disease. Neurobiol. Aging 52, 71–80 (2017).

29. Russo, N., Larson, C. & Kraus, N. Audio-vocal system regulation in children with autism spectrum disorders. Exp. Brain Res. 188, 111–24 (2008).

30. MacDonald, E. N., Johnson, E. K., Forsythe, J., Plante, P. & Munhall, K. G. Children's development of self-regulation in speech production. Curr. Biol. 22, 113–117 (2012).

31. Chen, X. et al. Sensorimotor control of vocal pitch production in Parkinson's disease. Brain Res. 1527, 99–107 (2013).

32. Mollaei, F., Shiller, D. M., Baum, S. R. & Gracco, V. L. Sensorimotor control of vocal pitch and formant frequencies in Parkinson's disease. Brain Res. 1646, 269–277 (2016).

